# The Atypical Fibrin Fibre Network in Rheumatoid Arthritis and its Relation to Autoimmunity, Inflammation and Thrombosis

**DOI:** 10.1101/2020.05.28.121301

**Authors:** Johannes A. Bezuidenhout, Chantelle Venter, Timothy Roberts, Gareth Tarr, Douglas B. Kell, Etheresia Pretorius

**Affiliations:** Department of Physiological Sciences, Faculty of Science, Stellenbosch University, Stellenbosch, Private Bag X1 Matieland, 7602, South Africa; Division of Rheumatology, Department of Medicine, Faculty of Medicine and Health Sciences, Stellenbosch University, PO Box 241 Cape Town, 8000, South Africa; Department of Biochemistry and Systems Biology, Institute of Systems, Molecular and Integrative Biology, Faculty of Health and Life Sciences, University of Liverpool, Crown St, Liverpool L69 7ZB, UK; The Novo Nordisk Foundation Centre for Biosustainability, Building 220, Chemitorvet 200, Technical University of Denmark, 2800 Kongens Lyngby, Denmark; University College London Hospital NHS Foundation Trust, 250 Euston Road, London, NW1 2PB.

**Keywords:** Rheumatoid Arthritis, Coagulation, Fibrinogen, Citrullination, CRP, SAA, ICAM-1, VCAM-1, Cardiovascular Risk

## Abstract

**Objective:** The risk of cardiovascular events in patients with RA is disproportionately heightened as a result of systemic inflammation. The relative effect of autoimmune-associated citrullination on the structure and thrombotic potential of fibrin(ogen) remains unknown. We therefore compared indices of vascular function, inflammation, coagulation and fibrin clot composition in RA patients with healthy controls and evaluated inter-parameter relationships.

**Methods:** Blood samples were collected from 30 RA patients and 25 age- and gender-matched healthy volunteers. Levels of SAA, CRP, ICAM-1 and VCAM-1 was measured using a sandwich immunoassay. Whole blood coagulation was assessed using Thromboelastography. Fibrin clot networks and fiber structure was investigated using Scanning Electron Microscopy. The detection and quantification of citrullination in formed fibrin clots were performed using a fluorescently labeled Citrulline monoclonal antibody with Confocal Microscopy.

**Results:** Concentrations of SAA, CRP and ICAM-1 were significantly elevated in RA patients compared to controls. TEG parameters relating to coagulation initiation (R and K), rate of fibrin cross-linking (α-Angle), and time to reach maximum thrombus generation (TMRTG) were attenuated in RA patients. Parameters relating to clot strength (MA, MRTG, TGG) did not statistically differ between RA and controls. Logistic regression modelling revealed stronger association between acute phase reactants (CRP, SAA) with TEG parameters than endothelial function markers. Microscopic analysis revealed denser networks of thicker fibrin fibers in RA patients compared to controls [median (interquartile range) 214 (170-285) *vs* 120 (100-144) nm respectively, p<0.0001, Odds ratio=22.7). Detection of multiple citrullinated regions within fibrin clot structures in RA patients, which was less prevalent in control samples (p<0.05, OR=2.2).

**Conclusion:** Patients with active RA display a coagulation profile that is dissimilar to general findings associated with other inflammatory conditions. The alteration of protein structures by autoimmune linked citrullination could play a role in determining the structure of fibrin and the potential of conferring a heightened thrombotic risk in RA patients.

## Introduction

Rheumatoid Arthritis (RA) is a chronic, systemic autoimmune disease characterized by both peripheral joint and extra-articular site inflammation, with an increased predisposition to a higher incidence of cardiovascular disease (CVD) (1, 2). CVD (including stroke and myocardial infarction) is almost 50% more common in RA patients than the general population and is the most frequent cause of early mortality (3). Traditional risk factors for CVD (age, hypertension, obesity, etc.), do not fully account for the elevated occurrence of CVD events, and thus RA (genetics and disease characteristics) has been identified as a strong independent risk factor (4). The interdependence of inflammatory and hemostatic pathways is well established and observable in multiple types of tissue, organs and pathologies (5). Disruption of the tightly regulated homeostatic control of immune and hemostatic systems could result in a rapid progression towards a prothrombotic tendency, a central cause of ischaemic stroke and myocardial infarction (6). This circumstance holds true for RA, with elevated levels of both pro-inflammatory [e.g. C-reactive protein (CRP) (7–11), Tumor necrosis factor alpha (TNFα) (7, 9, 11), Interleukin-6 (IL-6) (7–12), IL-1β (7) and Serum Amyloid A (SAA) (10, 13, 14)] and prothrombotic markers [e.g. D-dimer (8, 9, 11, 15, 16), Fibrinogen (10, 11, 16, 17), Tissue Factor (TF) (15), and von Willebrand factor (vWF) (8, 16, 18)], which is associated with one another (7, 19) and with the risk of future cardiovascular complications (20–23).

Key intermediaries of this manifestation are the structural components of formed thrombi. Soluble fibrinogen is cleaved by thrombin in order to form dense matrices of thin fibrous protein known as fibrin (24). Polymerized fibrin networks are essential for wound healing and other occlusive physiological processes (24). However, exposure to inflammatory biomarker stimuli [such as CRP (25), SAA (26), and pro-inflammatory cytokines (27, 28)] can result in the alteration of mechanical and viscoelastic properties of fibrin clots into a prothrombotic phenotype. This phenomenon has previously been observed in RA plasma clots (29, 30). Various immunopathogenic processes related to RA development can exert upstream amplification of the coagulation cascade as well as impairing fibrin clot dissolution (9, 19, 31).

Fibrin(ogen) is also a potent pro-inflammatory signaling entity itself, mainly through ligand-receptor interactions with immune cells that further propagates pro-inflammatory effects (32–36). The deimination of particular arginine residues in fibrin(ogen), known as citrullination, is a distinctive RA posttranslational modification that alters normal protein structure and function that confers antigenicity to modified proteins (37–43). The functional relationship between citrullination and the presence of a prothrombotic fibrin clot phenotype is still poorly understood. Some studies have shown that citrullination of fibrinogen prevents thrombin digestion and subsequent fibrinogenesis (44–46). However, the experimental conditions upon which these findings are based do not reflect physiological coagulation and is inconsistent with a predominantly hypercoagulable state seen in RA (47).

Inflammation-induced fibrin formation is equally present in RA synovial spaces as it is in circulation (Refer to **Figure 1**). Synovial coagulation is a key step in pannus formation, where fibrin provides the structural scaffold for immune cells that are responsible for synovial membrane disintegration and eventual joint damage (31, 48). Endothelial tissue dysfunction is a key process that facilitates this ubiquitous distribution of aberrant fibrin deposition in both synovia and vasculature. This pathophysiological state is characterized by the expression of cell adhesion molecules [intercellular cell adhesion molecule-1 (ICAM-1) and vascular cell adhesion molecule-1 (VCAM-1)], pro-inflammatory cytokines and pro-thrombotic markers (49, 50). This allows for the recruitment, translocation and propagation of inflammatory and thrombotic mediators across the synovial barrier (51–53).

**Figure 1:**
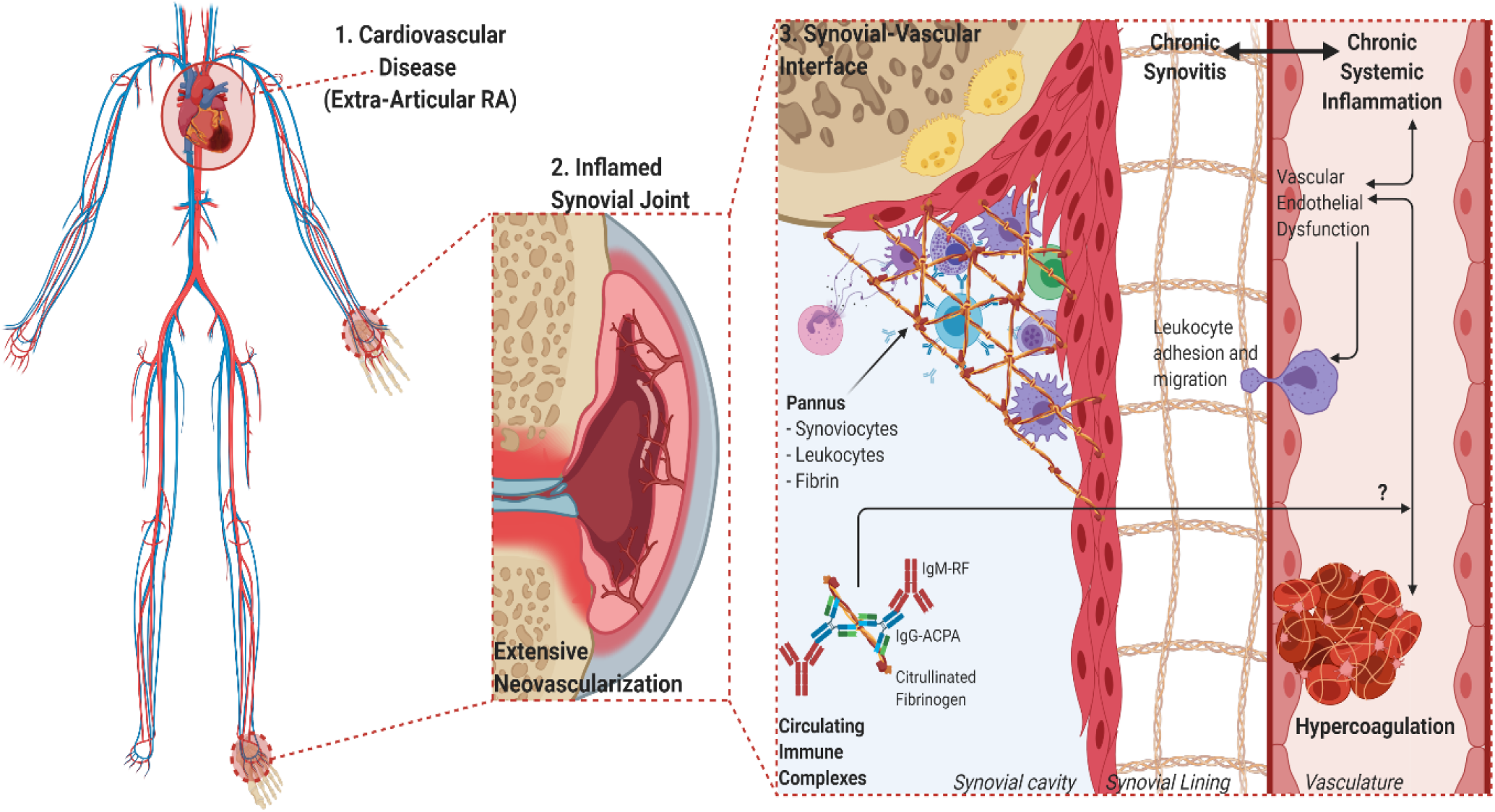
Overview of the overlapping processes of inflammation and coagulation in both synovial and vascular compartments. **1.** The chronic and systemic nature of the inflammatory response in RA characterizes the disease as an independent risk factor for CVD. **2.** The movement of leukocytes, inflammatory cytokines, procoagulant factors and immune complexes are aided by vascular endothelial dysfunction and neovascularization of hyperproliferative joint tissues. **3.**The role of fibrin(ogen) is integral to the formation of hyperplastic and destructive synovial tissue (pannus) and vascular thrombosis, while being a prominent self-protein target of aberrant citrullination and autoimmunogenicity in RA.

There is significant overlap in inflammatory pathways responsible for joint damage in RA and hypercoagulation, coupled with the fact that disease severity has been correlated to more adverse cardiovascular complications (21, 54, 55). It is therefore prudent that these processes and their relevant markers be examined systemically in RA, and not isolated to either vascular or synovial compartment. The aim of this study was to examine the extent to which the coagulation profiles and fibrin network architecture of RA patients are influenced by acute phase inflammation, endothelial dysfunction an autoimmune-related protein modification.

## Materials and methods

### Ethical considerations

Ethical approval for this study was given by the Health Research Ethics Committee (HREC) of Stellenbosch University (reference number: 6983). This study was carried out in strict adherence to the International Declaration of Helsinki, South African Guidelines for Good Clinical Practice and the South African Medical Research Council (SAMRC) Ethical Guidelines for research. Written consent was obtained from all participants (RA patients and healthy participants) prior to any sample collection.

### Study population

The RA sample group consisted of 30 patients (24 female and 6 male) that visited the Winelands Rheumatology Clinic (Stellenbosch, South Africa) for routine check-ups. All patients fulfilled the 2010 American College of Rheumatism/European League against Rheumatism (ACR/EULAR) classification criteria for RA diagnosis (56). The mean age of RA group was 53.4 years (range 22-75 years) with a mean disease duration of 10.5 years (range 1-39 years). RA participants were excluded from the study if they presented with other severe comorbidities (such as cancer or diabetes), existing cardiovascular disease or taking anticoagulant medication. RA participants were not excluded on the basis of any antirheumatic drug treatment or the use of glucocorticosteroids. The majority of RA patients (87%) were on a schedule of non-biologic disease modifying antirheumatic drugs (DMARDS, such as methotrexate, hydroxychloroquine, sulfasalazine, or leflunomide), while a lower proportion of patients were on biologic DMARDs (60%) and cortisone (14%, 5-10mg dosage). The control group consisted of 30 age- (mean: 53.9 years) and gender- (22 female and 8 male) matched volunteer blood donors. The inclusion criteria for healthy controls were: (i) no history of thrombotic disease or inflammatory conditions (ii) no use of any chronic medication (ii) no use of anticoagulant therapy (iii) non-smokers (iv) females not taking contraceptive medication or hormone replacement therapy (v) females that are not pregnant or lactating. All demographic information is summarized in **Table 1**.

**Table 1:** Demographic and clinical characteristics of study participants.

| Risk Factor | RA (n=30) | Control (n=30) |
| --- | --- | --- |
| Age (years) | 53.5 (22-75) | 50 (28-79) |
| Women [n (%)] | 24 (80%) | 22 (73.3%) |
| Smokers [n (%)] | 3 (10%) | - |
| Disease duration (years) | 6.5 (1-39) | - |
| Hypertension [n (%)] | 5 (16.7%) | - |
| Hypercholesterolemia [n (%)] | 3 (10%) | - |
| Disease activity (DAS-28) classification [n (%)] |  | - |
| - Remission | 1 (3.3%) |  |
| - Low | 17 (56.7%) |  |
| - Moderate | 4 (13.3%) |  |
| - High | 8 (26.7%) |  |
| Autoantibody seropositivity [n (%)] |  |  |
| - RF-/Anti-CCP<sup>-</sup> | 1 (3.3%) |  |
| - RF<sup>+</sup>/Anti-CCP<sup>-</sup> | 6 (20%) |  |
| - RF-/Anti-CCP<sup>+</sup> | - |  |
| - RF<sup>+</sup>/Anti-CCP<sup>+</sup> | 23 (76.7%) |  |
| Conventional DMARD use [n (%)] | 26 (86.7%) | - |
| Biologic DMARD use [n (%)] | 18 (60%) | - |
| Cortisone use [n (%)] | 13 (43%) | - |

Values expressed as median and interquartile range (IQR) unless otherwise stated. Disease activity scores (DAS-28), were categorized in RA patients as remission, low, moderate or high (59). For full demographic information and specified list of disease modifying anti-rheumatic drugs (DMARDs) please refer to full study database in supplementary material.

### Blood sampling

Whole blood (WB) samples were collected in vacutainer tubes using 3.8% sodium citrate as anticoagulant. Blood drawing on all participants was performed by a qualified nurse, or phlebotomist by sterile puncture of the antecubital vein. Blood tubes were incubated at room temperature for a minimum duration of 30 minutes prior to the commencement of any whole blood analysis. In order to obtain platelet poor plasma (PPP), sodium citrated blood tubes were centrifuged at 3000x*g* for 15 minutes, aliquoted into Eppendorf tubes and stored at −80°C until further analysis.

### Thromboelastography^®^

Analysis of dynamic coagulation kinetics were performed on RA and control WB by means of Thromboelastograph^®^ (TEG^®^) 5000 Haemostasis Analyzer System (Haemonetics^®^, 07-033). In brief, coagulation is initiated by recalcification of 340μL WB with 20μL of 0.2mM Calcium chloride (CaCl_2_) (Haemonetics^®^, 7003) in a disposable TEG^®^ cup (Haemonetics^®^, 6211). Various kinetic clotting parameters are determined by assessing the resistance that the forming thrombus provides against the oscillating pin of the instrument (measuring at 37°C). Parameters derived from the thromboelastograph tracing consist of: reaction time (R, time from test start to initial fibrin formation in minutes), kinetics [K, time required to reach an amplitude (clot thickness) of 20mm, in minutes], alpha angle (α, rate of fibrin cross linking indicated by degrees), maximal amplitude (MA, maximum strength of formed clot in millimeters), maximum rate of thrombus generation (MRTG, in dynes.cm^−2^.s^−1^), time to maximum rate of thrombus generation (TMRTG, in minutes) and total thrombus generation (TTG, in dynes.cm^−2^).(57)

### Scanning electron microscopy

The ultrastructure of fibrin networks and individual fibrin fibers were examined using scanning electron microscopy (SEM). In summary, clots were prepared from thawed PPP samples of RA patients (n=10) and controls (n=10) by addition of 5μL human thrombin (provided by South African National Blood Service) to 10μL PPP on a glass coverslip and transferred to a 24-well plate. Preparation consisted of washing with 10X Gibco^®^ phosphate-buffered saline (PBS, pH 7.4) (ThermoFisher Scientific, 10010015), chemical fixation with 4% Paraformaldehyde (PFA) (Sigma-Aldrich, P6148) and then 1% Osmium Tetrahydroxide (OsO_4_) (Sigma-Aldrich, 75632), followed by dehydration with increasing grades of ethanol and 99.9% Hexamethyldisilizane (HMDS) (Sigma-Aldrich, 37921) [for detailed protocols please refer to (57)]. Samples were carbon coated using a Quorom Q150T E carbon coater. Images were captured at an electron high tension (EHT) of 1kV using a high resolution InLens detector of the Zeiss Merlin™ (Gemini II) FE SEM (Carl Zeiss Microscopy, Munich, Germany). Fibrin fiber diameters representative of each respective sample group (RA and control) was determined by means of image analysis software ImageJ (Version 1.52p). Three representative micrographs (78,98μm^2^ image size, 10 000x magnification) were calibrated to scale and overlaid with a non-destructive grid (2μm^2^ tile size). Single representative fibrin fibers were measured in 28 tiles per image, producing 84 fiber diameter measurements per sample (method illustrated in **figure 2**).

**Figure 2:**
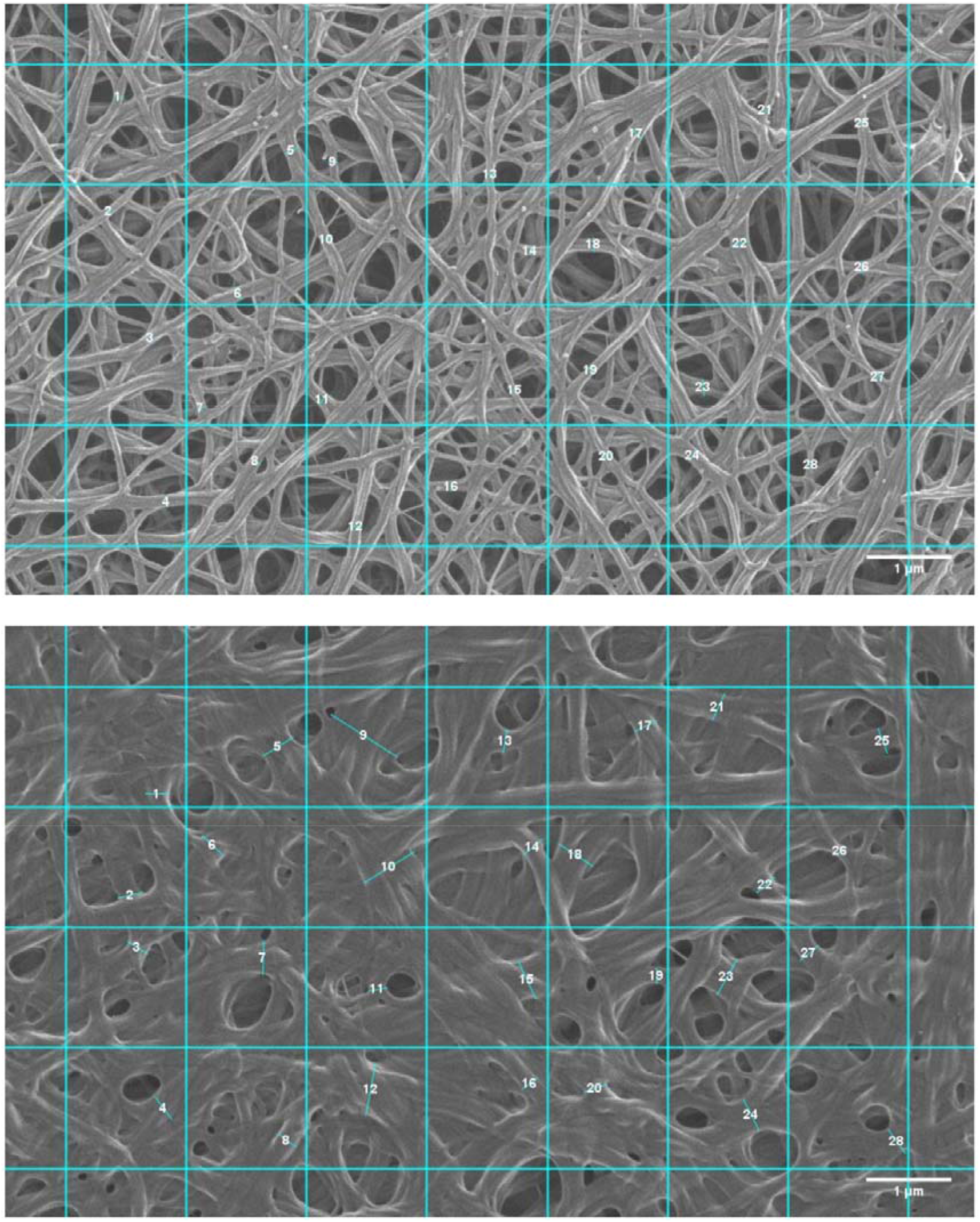
Fibrin fiber diameter measurement scheme. SEM micrographs of a prepared PPP clot from a representative healthy control (top) and RA (bottom).

### Vascular injury panel analysis

Plasma concentrations of soluble ICAM-1, VCAM-1, CRP, and SAA were measured by sandwich immunoassay [Meso Scale Discover (MSD) Vascular Injury Panel (V-plex) 2 (human) kits, catalogue number K15198D]. RA (n=30) and control (n=30) PPP samples and reagents were prepared as per manufacturer’s protocol. Samples were run in duplicate and measurements read on an MSD Discovery Workbench 4. Analyte concentrations were calculated from the calibration curve generated by absorbance measurements of manufacturer supplied calibrator standards.

### Confocal laser scanning microscopy

In order to determine the extent of protein deamination in fibrin networks, PPP aliquots of RA samples (n=10) and control samples (n=10) were thawed and fibrin clots prepared (refer to SEM method) on glass microscope slides in a dark room. Samples were fixed with 4% PFA, washed 3x with PBS, and blocked with 5% Goat serum solution (Abcam, ab7481) for 30 minutes. Clots were then stained with a 1:50 dilution Citrulline Monoclonal Antibody (2D3.1) (Thermo Fisher Scientific, Cat #MA5-27573) and incubated for 1 hour. Following another 3x PBS wash to remove unbound antibodies, samples were then stained with 1:200 dilution Goat Anti-Mouse IgG Secondary antibody conjugated to AlexaFluor 488 (Thermo Fisher Scientific, Cat #A327273) and incubated for 1 hour. Slides were washed 3x with PBS to remove unbound antibody, allowed to dry, and mounted with a glass coverslip. Samples were viewed with a Zeiss LSM 780 Confocal laser scanning microscope (CLSM) with a Plan-Apochromat 63x/1.4 oil DIC M27 objective. AlexaFluor488 was excited with the 488nm laser and emission was detected at 508-570nm. Three representative micrographs per sample were analysed for fluorescent particle distribution using ImageJ (Version 1.52p). Images were calibrated to scale, and a global threshold (27 pixel cut-off) applied to all analysed micrographs.

### Statistical analysis

Statistical analysis was performed using R version 4.0. Specifically, univariate logistic regression was performed to determine odds ratios (OR) for experimental variables using the logistic model in the rstanarm package (with default priors). ORs and 95% confidence intervals were extracted in the corresponding unit system (i.e. not z-scaled) for all variables except Fibrin fiber diameter and citrulline particle number shown in Table 4 which *are* z-scaled to aid interpretation. Tables 2 and 3 show ORs after adjustment for age and gender with unadjusted analysis identifying the same significance and effect sizes. In addition, results for classical statistical tests are reported as follows. The distribution of sample datasets for each variable experimental was determined using the Shapiro-Wilk test. Accordingly, *p*-values for each variable comparing RA to healthy controls were calculated using either a Mann-Whitney *U* test for nonparametric data or a Student t-test for parametric data. Statistical significance was set at *p*<0.05. One can see close alignment between all these and the OR results.

**Table 2:** Vascular injury panel (V-Plex) analysis.

| V-Plex Analyte | RA<br>(n=30) | Control<br>(n=25) | Adjusted OR<br>(95% CI) | P-value |
| --- | --- | --- | --- | --- |
| CRP<br>(µg/mL) | 4.25<br>[1.89-8.11] | 1.26<br>[0.43-3.15] | 1.29<br>(1.09-1.62) * | 0.0011** |
| SAA<br>(µg/mL) | 4.98<br>[2.56-8.84] | 1.52<br>[0.64-2.24] | 1.25<br>(1.08-1.55) * | <0.0001**** |
| sICAM-1<br>(pg/mL) | 377.82<br>[295.69-460.49] | 289.7<br>[225.34-365.56] | 1.008<br>(1.0021-1.014) * | 0.0202* |
| sVCAM-1<br>(pg/mL) | 359.83<br>[283.01-394.20] | 325.8<br>[253.73-422.62] | 1.001<br>(0.995-1.007) | 0.9 |

Analyte concentrations for RA and control are expressed as median [IQR]. Odds ratios for age- and gender-adjusted logistic regression model is listed with the 95% confidence interval, while level of statistical significance was set at $p<0.05$ for calculated Mann-Whitney test values. \*Denotes statistically significant differences in variables between RA and controls.

**Table 3:** Thromboelastography^®^ analysis.

| TEG® Parameter | RA (n=30) | Control (n=30) | Adjusted OR (95% CI) | P-value |
| --- | --- | --- | --- | --- |
| R (min) | 8.3 [7.5-9.5] | 11.9 [8.3-15.1] | 0.675 (0.528-0.827) * | 0.0007*** |
| K (min) | 2.6 [2.2-3.2] | 3.6 [2.6-4.5] | 0.539 (0.308-0.847) * | 0.0196* |
| $\alpha$ (°) | 63.2 [57.6-68.6] | 54.2 [48.4-60.5] | 1.15 (1.07-1.27) * | 0.0006*** |
| MA (mm) | 55.9 [52.0-61.7] | 58.9 [54.5-63.9] | 0.960 (0.889-1.03) | 0.1646 |
| MRTG (dyn·cm<sup>-2</sup>·s<sup>-1</sup>) | 5.01 [3.89-6.02] | 4.23 [3.26-5.85] | 1.03 (0.767-1.40) | 0.2745 |
| TMRTG (min) | 11.67 [9.73-12.96] | 17.43 [12.48-21.13] | 0.737 (0.607-0.858) * | 0.0001*** |
| TTG (dyn·cm<sup>-2</sup>) | 633 [544.1-775.2] | 719.3 [602.1-879.8] | 0.997 (0.995-1.00) | 0.0899 |

Viscoelastic coagulation measurements for RA and control are expressed as median [IQR]. Odds ratios for age- and gender-adjusted logistic regression model is listed with the 95% confidence interval, while level of statistical significance was set at $p < 0.05$ for calculated Mann-Whitney test values. \*Denotes statistically significant differences in variables between RA and controls.

**Table 4:** Microscopy Image Analysis.

| Imaging Parameter | RA (n=10) | Control (n=10) | OR (95% CI) | P-value |
| --- | --- | --- | --- | --- |
| Fibrin fiber diameter (nm) | 214 (170-285) | 120 (100-144) | 22.732 (17.085-31.441) | <0.0001 **** |
| Fluorescent citrulline particles | 3279 (1966-4260) | 1095 (411-2937) | 2.2268 (1.2238-4.3527) | 0.0355 * |

Evaluation of plasma clots using image analysis software (ImageJ) based techniques for representative RA (n=10) and Healthy control (n=10) subjects. Fibrin fiber diameters were determined from SEM micrographs, while fluorescent particle analysis was calculated from CLSM micrographs. Values are expressed as median [interquartile range]. Scaled odds ratios for the logistic regression model are listed with their 95% confidence intervals, while level of statistical significance was set at $p < 0.05$ for calculated Mann-Whitney test values. \*Denotes statistically significant differences in variables between RA and controls.

## Results

### Subjects

Demographic information of all study participants is listed in **Table 1**. The RA sample group closely resembles the general population distribution for age (median: 54 years) and sex (80% female) of the disease (58). The control group of healthy volunteers was closely matched to the RA group with regards to age (median: 50 years) and sex (73% female). The RA sample group was heterogeneous with respect to clinical presentation, with most patients on an anti-rheumatic drug therapy regime. The majority of RA patients also presented with positive titers for anti-cyclic citrullinated peptide (CCP) (77%) and rheumatoid factor (97%) autoantibodies.

### Confirmation of altered inflammatory and vascular function profile in RA

Circulating concentrations of endothelial function and acute phase markers are shown in **Table 2**. Previous studies have shown these markers (CRP, SAA, sVCAM-1 and sICAM-1) to be associated with a prothrombotic state and increased CVD risk in RA(7, 13, 60-62), and was therefore measured in this study to determine the extent to which systemic inflammation influences viscoelastic and structural clot properties. As expected, all markers were elevated in RA compared to controls [CRP (median 4.25 μg/mL vs 1.26, OR=1.29), SAA (4.98 μg/mL vs 1.52, OR=1.29), sICAM-1 (378.82 ng/mL vs 289.70, OR=1.01), sVCAM-1 (359.83 ng/mL vs 325.80, OR=1.00)].

### Functional coagulation assessment indicates a prothrombotic tendency in RA

Whole blood coagulation parameters as measured by TEG^®^ are listed in **Table 3**. Limited viscoelastic assessment of coagulation in RA has been performed to date (60, 63, 64), with TEG^®^ not commonly used in rheumatology practice (65–67). RA patients showed significantly altered rates of clot formation compared to healthy controls. This included shortened clot initiation (R; OR=0.675, p=0.0007 and K; OR=0.539, p=0.0196), augmented fibrin cross-linking (α; OR=1.15, p=0.0006) and shortened time to maximal thrombus formation (TMRTG; OR=0.737, p=0.0001). Measures of overall clot strength (MA) and growth (TTG) were attenuated in RA but did not statistically differ from those of controls.

### Association of vascular injury biomarkers to thrombotic parameters indices

Significantly altered inflammatory and coagulation indices in RA, as determined by an adjusted logistic regression model, are represented by box-and-whisker plots (**Figure 3**). Additionally, the distribution of RA and control experimental values are plotted and arranged in a lattice (**Figure 4**) to exhibit correlation values between intra- and inter-assay variables. CRP shows positive and significant association with SAA (*r* = 0.6) and sICAM-1 (*r* = 0.52). Further strong correlation existed between respective endothelial markers (sICAM-1 and sVCAM-1, *r*=0.62), and between all TEG^®^ parameters (R, K, α, TMRTG). CRP and SAA display stronger association with thromboelastographic parameters than the cell adhesion molecules.

**Figure 3:**
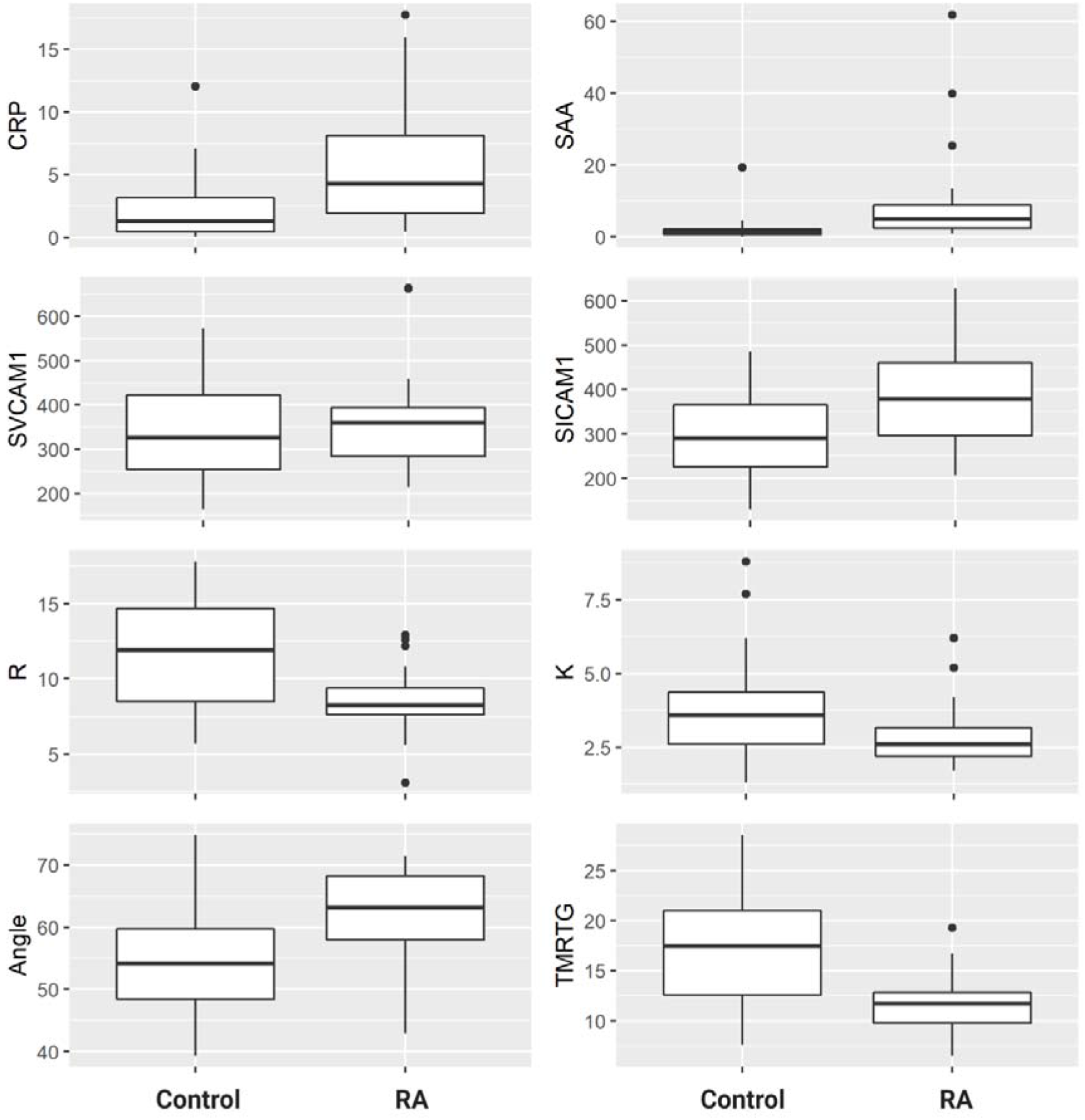
Box-and-whisker plots for statistically significant TEG^®^ and V-plex parameters. Boxes represent the median and IQR. Whiskers indicate upper (75^th^ percentile + 1.5*IQR) and lower (25^th^ percentile −1.5*IQR) extremes. Outlier values are indicated by black dots

**Figure 4:**
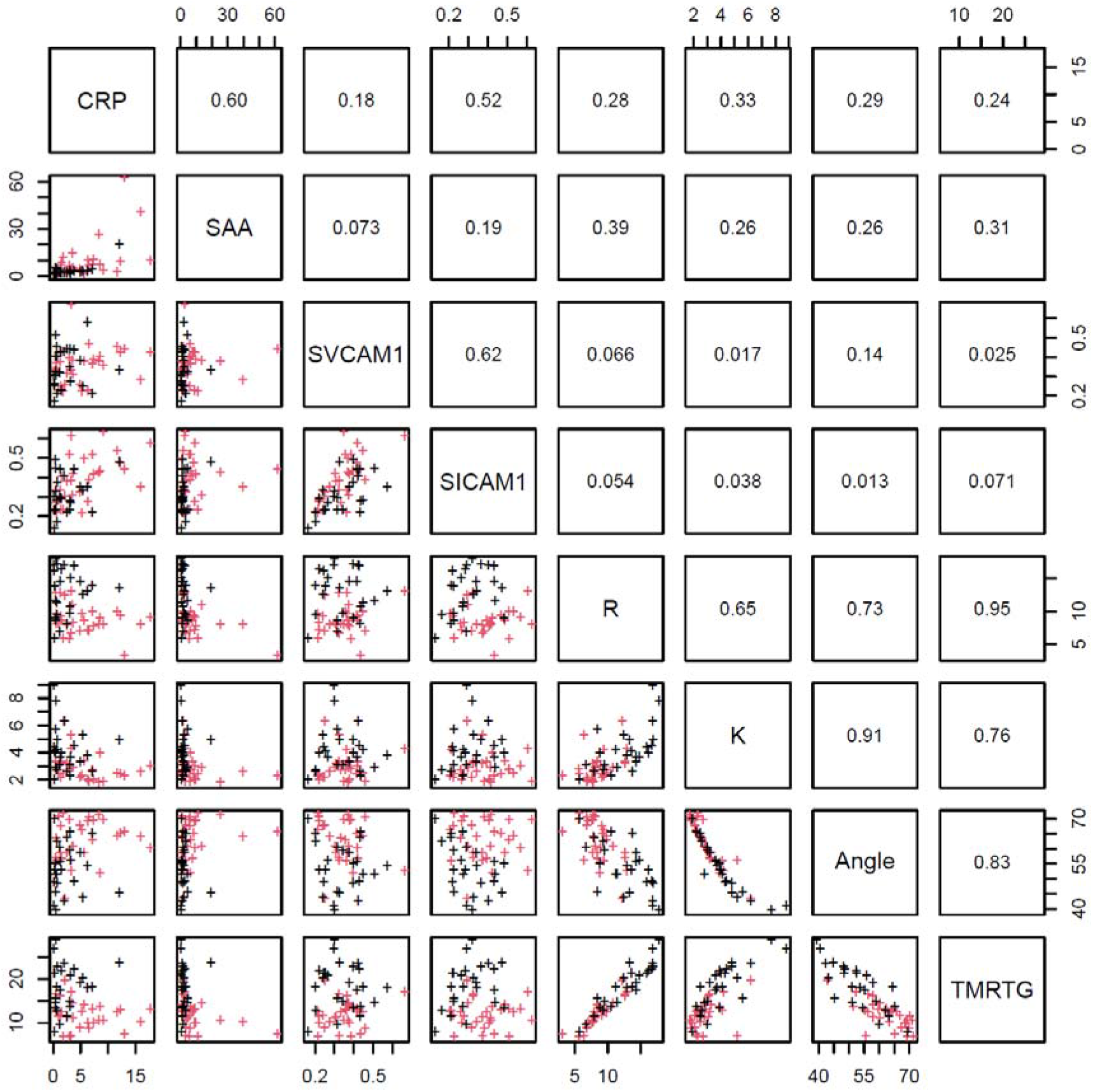
Lattice of cross-plots and correlation values across statistically significant TEG^®^ and V-Plex measurements. The sample values are indicated by crosses (RA: red, Control: black). Strong intra-analysis correlation exists between CRP with SAA and sICAM-1, between sICAM-1 and VCAM-1, and between all TEG parameters (R, K, Angle, TMRTG). CRP and SAA show moderate correlation with inflammatory and endothelial markers.

### SEM analysis exposes anomalous fibrin network architecture in formed RA clots

Further investigation into the apparent modification of the clot structure in RA was carried out by means of SEM. **Figure 2** illustrates the scheme followed for the measurement of fibrin fiber diameters in PPP clots of RA (n=10) and Control (n=10) samples. Results (refer to **Table 4**) indicate that fibrin fiber diameters in representative areas were significantly increased in RA versus controls (median: 214nm vs 120nm, OR=22.732). Examining the networks qualitatively, it is evident that *ex vivo* formed clots from RA samples have denser, less porous fibrin networks. **Figure 5 B-D,** representative of the RA sample group, illustrates the amalgamation of fibrin monomers that contribute to increased fibrin fiber diameter and overall network density. This contrasts sharply with the ultrastructural attributes of **Figure 5A** (Healthy control sample), which demonstrates thinner protein strands and a more permeable fibrin network.

**Figure 5:**
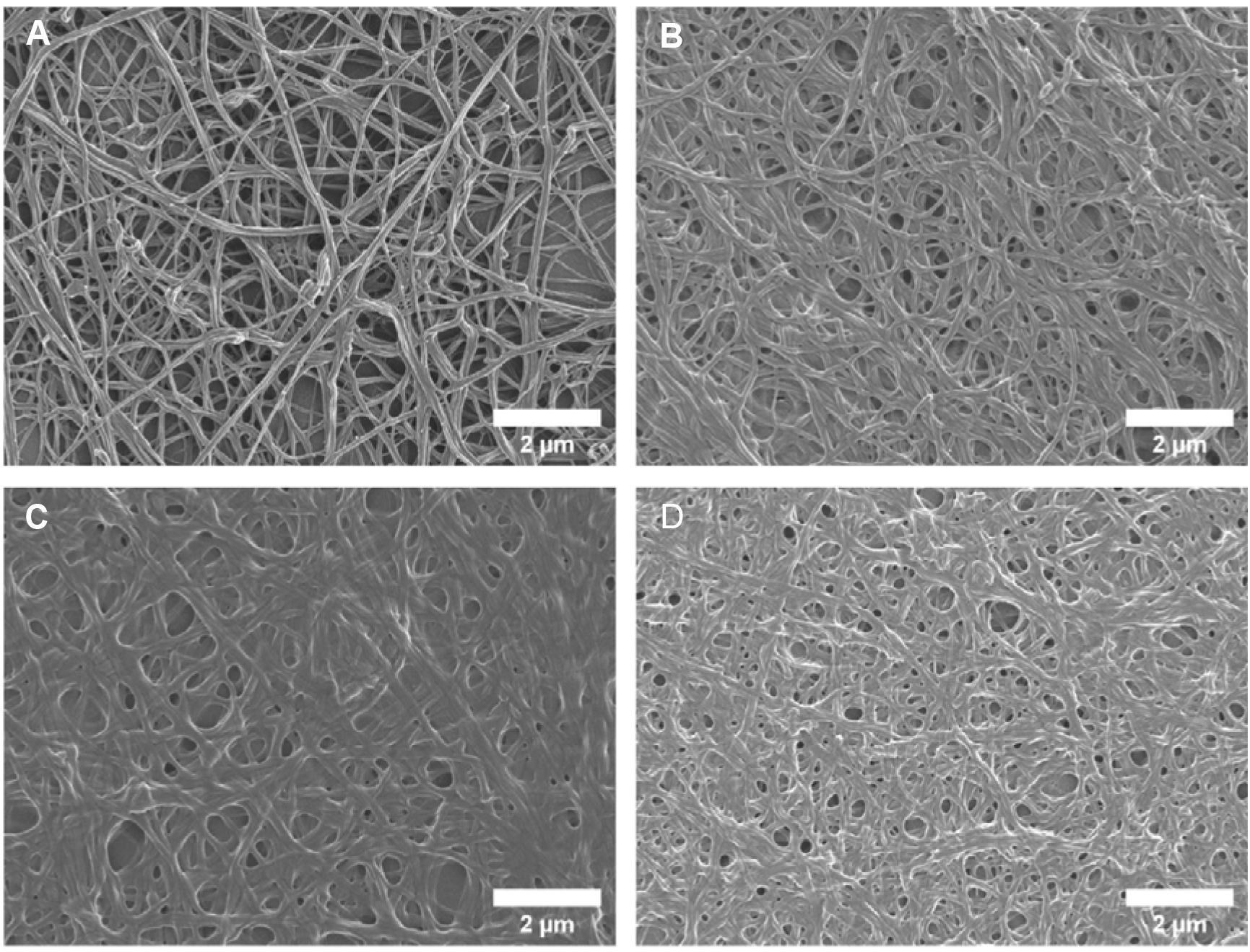
Scanning electron micrographs of the fibrin network ultrastructure. Representative micrographs of the fibrin network in healthy controls (A) and RA patients (B-D). The altered clot ultrastructure in RA, consisting of less permeable networks of thicker fibrin fibers, represents a prothrombotic phenotype.

### Confocal analysis of plasma clots reveals a higher number of citrullinated sites in RA fibrin networks compared to controls

Additionally, we probed whether autoimmune-related modifications of coagulation proteins are distinctive to RA patients compared to healthy individuals. Fibrin and fibrinogen are well known extracellular targets for peptidylarginine deiminase (PAD) enzymes (37, 68), with citrullinated isoforms eliciting the generation of anti-CCP antibodies in large proportions of RA patients (40, 41). Citrullinated fibrin deposits are common synovial compartment manifestations, but this has not been investigated in plasma. The determination of the net-effect of coagulation factor citrullination on hemostatic outcome also remains elusive. To investigate the presence and extent of potential citrullination in RA (n=10) and Control (n=10) PPP thrombi, fluorescence analysis using a Citrulline-identifying monoclonal antibody with CLSM was performed (**Figure 6**). Acquired image data (**Figure 7**) suggests a relatively higher proportion of citrulline particles in RA fibrin networks versus controls (OR=2.2268).

**Figure 6:**
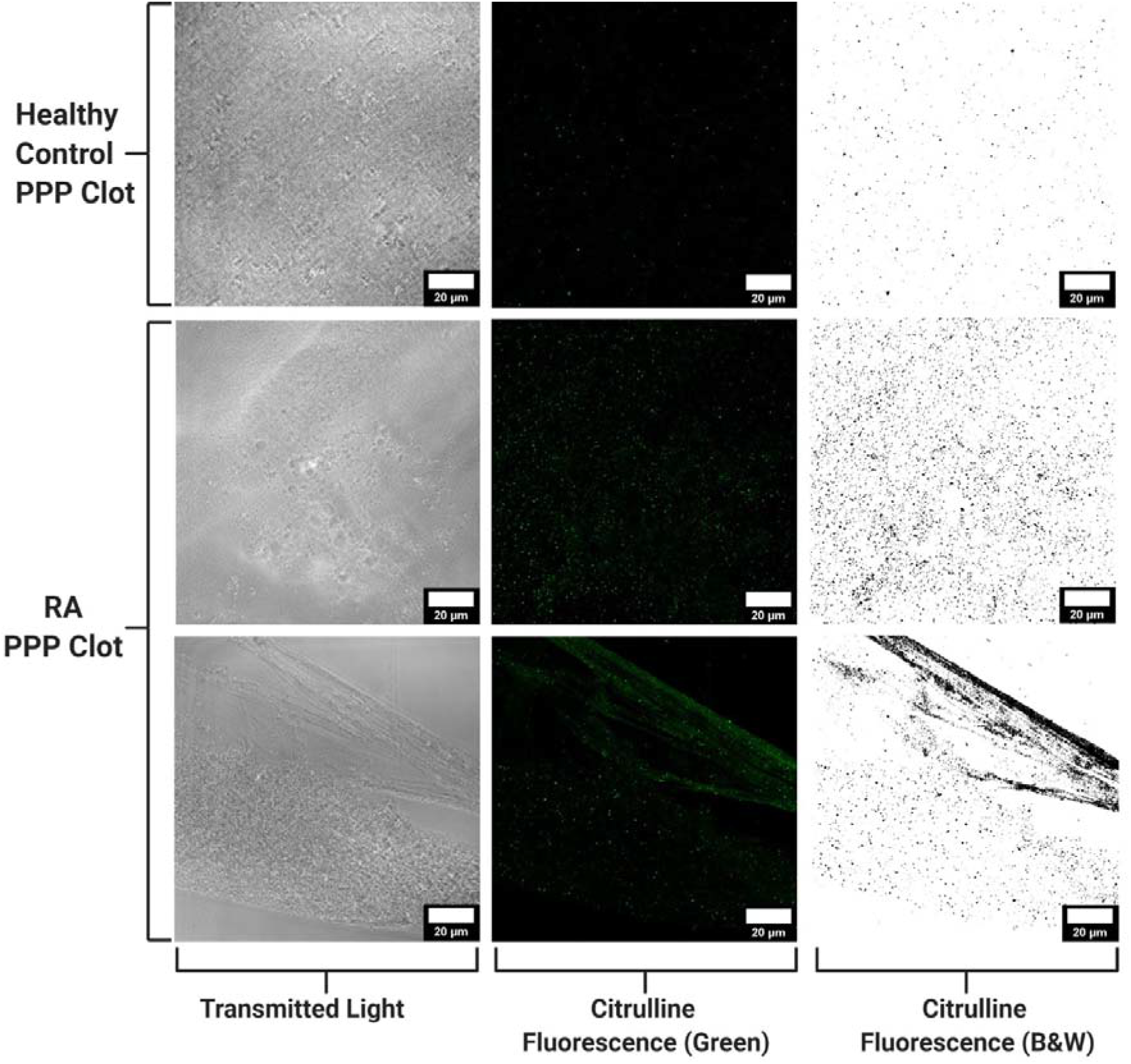
Confocal microscopy of PPP clots stained with a Citrulline monoclonal antibody. Confocal micrographs of representative control and RA samples. Each row represents identical areas captured of fibrin clot preparations using transmitted light to illustrate fibrin network topography (left-hand column) and a green channel (middle column) to identify fluorescent particles corresponding to citrulline residues. Fluorescent images are also provided in binary (right-hand column) to better illustrate differences seen between RA and control samples with respect to citrulline-coupled fluorescence. Particle analysis confirmed the observable presence of enhanced fluorescent signal in RA samples (n=10) versus healthy controls (n=10).

**Figure 7:**
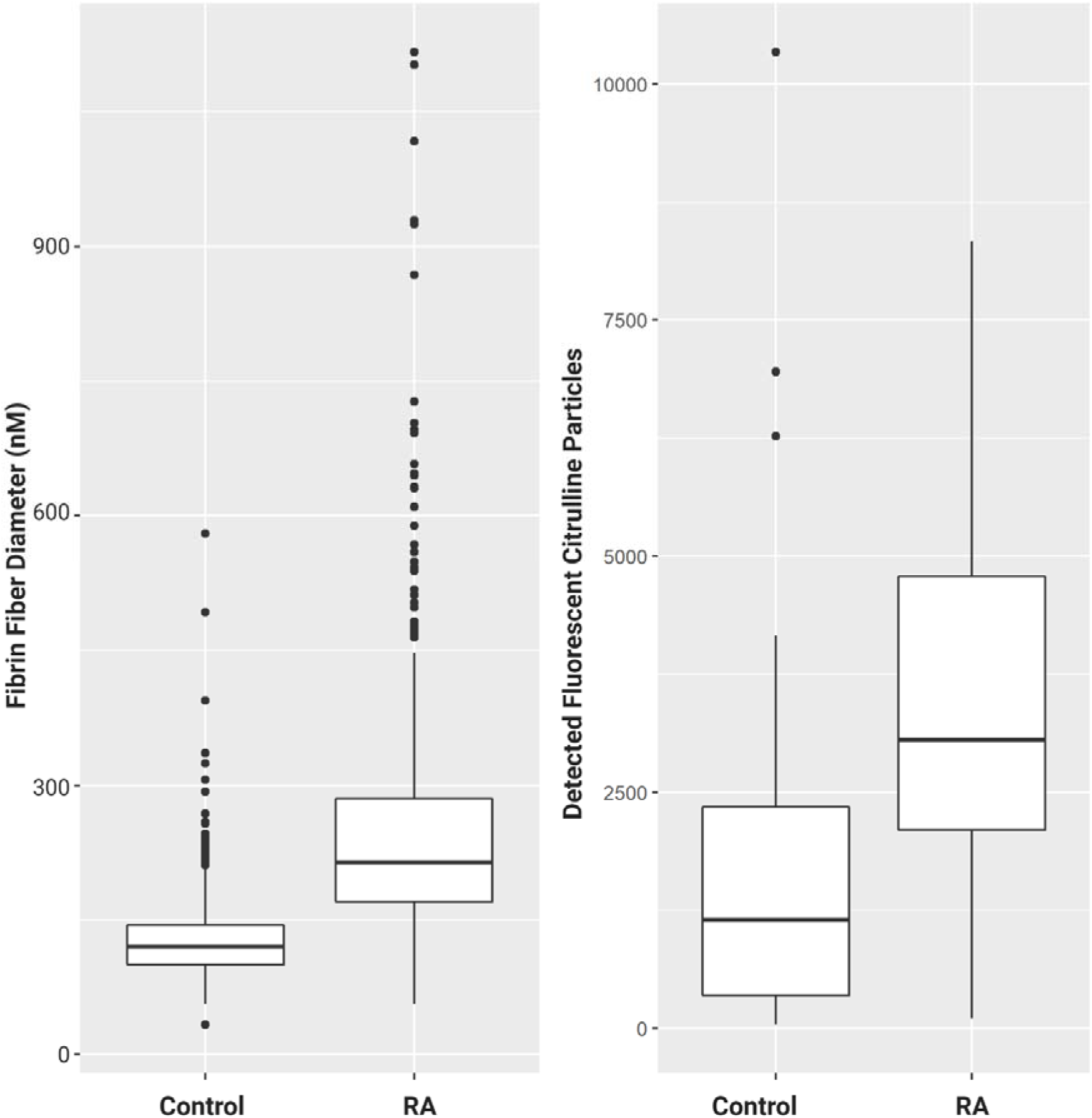
Box-and-whisker plots for micrographs analysis. Outlier values are indicated by black dots. In comparison to healthy individuals, RA subjects exhibited significantly increased fibrin fiber diameters (median: 214nm vs 120nm, OR=22.7) and detected fluorescent areas (median: 3279 vs 1095, OR=2.23) of fibrin citrullination.

## Discussion

There is a need to bridge translational gaps between RA immunopathogenesis and systemic vascular and hemostatic irregularities. The development of RA auto-immunogenicity precedes the onset of joint disease (69, 70). The link between RA autoimmune patterns and its possible role in exacerbating thrombosis is still poorly understood. Crosstalk between immune and hemostatic systems with the endothelium represents a critical interface in which both arthritic and cardiovascular pathologies are initiated and propagated. We therefore analyzed a panel of biomarkers that are representative of this dynamic milieu, is associated with RA disease severity and CVD.

Levels of both acute phase reactants (CRP and SAA) were significantly elevated in RA patients (**Table 2**) and showed a strong association (**Figure 4**). This was expected as acute phase reactant concentrations rise dramatically under acute inflammatory states, with both CRP and SAA shown to reliably predict disease severity and CVD risk in RA (13, 14, 22, 71, 72). CRP can bind to surface receptors on monocytes, endothelial cells and platelets thereby altering their function to propagate hypercoagulable conditions (73, 74). The ability of CRP and SAA to induce TF expression has been demonstrated in various cell types (13, 14, 22, 71, 72, 75–81) and by *in vivo* studies (82, 83). Additionally, acute phase reactants can also suppress fibrinolysis by promoting expression of plasminogen activator inbibitor-1 (PAI-1) (82) and inhibiting TF pathway inhibitor (TFPI) expression (81, 83). CRP is also able to activate endothelial cells and cause the expression of cell adhesion molecules (ICAM-1 and VCAM-1) (84, 85). It should however be noted that cellular effects of CRP have been disputed as being primarily caused by bacterial contaminants in CRP preparations rather than the protein itself (86). SAA is the precursor to amyloid A (AA) protein, which form insoluble fibrillar depositions in major organs as a result of long-term inflammation (87). RA has been prominently implicated as a pre-existing condition for the development of potentially fatal AA amyloidosis (88, 89). Known for being primarily hepatically synthesized, synovial tissue cells (90–93) and chondrocytes (94) can be articular sources of SAA production. Within the synovium SAA promotes pro-arthritic processes through the expression of cell adhesion molecules (95, 96), cytokines (96), and matrix degrading enzymes (92, 95) by local tissue. SAA showed slightly weaker association with sICAM-1 compared with CRP in our studies (**Figure 4**).

Levels of soluble cell adhesion molecules (sICAM-1 and sVCAM-1) are the most accessible form in which to determine endothelial activity. They are also strong predictive biomarkers for CVD in RA (61, 62, 97). Increased levels of CAMs indicate endothelial dysfunction that facilitates pro-inflammatory and prothrombotic conditions (7). ICAM-1 is a prominent receptor for fibrinogen (98), with interaction fortifying endothelial adhesion and migration of leukocytes (99–101), endothelial-platelet adhesion (102–104), and causes vasoconstriction (105, 106). ICAM-1 signaling pathways can also promote endothelial tissue factor expression (107). The role of VCAM-1 in directly promoting coagulation is not as well understood, but does contribute to atherosclerotic plaque formation as a result of adhering to PBMCs (108, 109) sICAM-1 and sVCAM-1 concentrations were elevated in RA (**Table 2**) but were not as strongly associated with coagulation function as CRP and SAA (**Figure 4**). This belies the fact that CAMs present on cell surfaces play more facilitative roles in thrombotic diseases rather than instigating them.

After confirming the presence of a systemic inflammatory state in RA patients, we investigated the possible repercussions thereof on the coagulation profile of study participants. Thromboelastography^®^ is a hemostatic function test that measures the rate, strength, and stability of clot formation through the viscoelastic changes induced by fibrin polymerization. Despite its widespread clinical use, especially as point-of-care instruments in surgical settings, TEG^®^ is not often utilized in Rheumatology practice. The advantage that TEG^®^ provides over conventional coagulation tests [such as prothrombin time (PT), activated partial thromboplastin time (aPTT) and D-dimer] is that it measures global hemostatic function and outcome rather than single time points or pathways in the coagulation cascade (110). Türk *et al.* (2018) is the only recent study that has assessed thrombotic tendency in RA patient with thromboelastographic assessment (60) Using a modified version of TEG^®^, known as rotational thromboelastometry (ROTEM), they found that RA disease markers (CRP, DAS-28) correlated strongly with ROTEM parameters. Our findings show that coagulation initiation was amplified in RA patients with shortened velocity parameters of clot formation (R, K, α, TMRTG) (**Table 3**) and these indices were moderately associated with levels of SAA and CRP (**Figure 3**).

Parameters relating to clot strength (MA, TTG) were attenuated in the RA sample group, but did not statistically differ from healthy controls (**Table 3**). Thus, although the blood clots form rapidly it leads to a weak clot. The mechanics of clot formation as measured by TEG^®^ have been related to the risk of adverse ischaemic events (111). Excessive hepatic production of fibrinogen is highly prevalent in RA (10). Increased plasma fibrinogen concentration is a strong contributing factor to hypercoagulation (112). Fibrin(ogen) is susceptible to structural and functional modifications by certain inflammatory molecules, including CRP (25), SAA (26), and certain bacterial virulence factors (113–115). Fibrin(ogen) is also prone to post-translational modification that relates to the generation of auto-immunogenicity in RA – the relevance of this process was investigated and is discussed below.

Evaluating fibrin gel matrices visually can reveal much about thrombotic potential under inflammatory conditions. Denser fibrin fiber networks are accompanied with increased resistance to fibrinolysis and is associated with the risk for thrombotic events (reviewed by Undas and Ariëns, 2011) (116). These structural properties can be viewed and functionally assessed with a high degree of resolution using SEM (57) Our analysis revealed denser fibrin networks in RA prepared *ex vivo* PPP clots compared to controls (**Figures 2 and 5**). This is consistent with a prothrombotic phenotype observed in previous studies that have inspected the fibrin network in RA (29, 30). Furthermore, we determined a relative measure of fibrin fiber diameter using an image analysis software-based technique (**Figure 2**). The average diameter of fibrin fibers was larger in RA clots compared to controls (**Figure 7**). This can be clearly seen in the presented micrographs, were there is an apparent amalgamation of single fibrin fibers in RA preparations (**Figures 2 and 5).** The appearance of very thin fibrin strands (approximately 100nm in diameter) was not readily detectable in the RA fibrin networks. Some studies have indicated that thin fibrin fibers have higher tensile strength than thicker fibers, concluding that dense networks consisting of predominantly thin fibers are more resistant to degradation (117, 118). Fibrin networks of this nature in RA were observed by Vrancic *et al.* (2019) (119). However, study by Buclay *et al.* (2015) concluded that thicker fibers are more resistant to plasmin degradation than thinner fibers, owing to their ability to elongate during lysis (120). It is apparent that our investigation into the structural properties of fibrin networks in RA and its relation to hemostatic function has a rather deceptive appearance. Our group has previously shown that fibrin networks in multiple inflammatory conditions (113, 121–123) and in plasma exposed to inflammatory stimuli (26, 114, 124) stain positive for amyloid-specific dyes. This constitutes the appearance of misfolded, protein aggregates with enriched β-sheet content that confers a higher degree of insolubility (125). It may therefore be possible that the fibrin network in RA may undergo a similar transition. This, coupled with impairment of fibrinolytic pathways, presents a possible explanation for high thrombotic risk seen in RA patients.

Distinct protein modifications related to the generation of autoimmunity in RA could present an additional complication when attempting to expound underlying mechanisms responsible for excessive thrombotic risk. Citrullination is a post-translational modification in which positively charged arginine are deiminated by peptidylarginine deiminase (PAD) enzymes to form neutrally charged citrulline (126). PADs are usually localized to intracellular environments and requires calcium for catalysis (127). PADs become active under inflammatory and apoptotic conditions where enzymes are liberated to extracellular spaces and exposed to sufficient calcium concentrations for catalytic function (126). Two isoforms of PAD (PAD2 and PAD4) are primarily responsible for generating citrullinated antigens that incite the generation of anti-citrullinated protein antibodies (ACPAs), a hallmark serological feature of RA (128). Fibrinogen and fibrin are prominent substrates for PAD2 and PAD4 and autoantibodies targeting citrullinated fibrin(ogen) have been identified (40, 43, 68, 129–131). The pathogenicity of citrullinated fibrin(ogen) immune complexes have been demonstrated both *in vitro(33)* and *in vivo(132, 133)*. Citrullinated fibrin deposits are also common manifestations within synovial cavities, where it contributes to self-perpetuating inflammatory processes (48, 134). Our findings provide novel evidence for the citrullination of fibrin within vasculature which is more prominent in RA plasma compared to control plasma (**Figures 6 and 7**). Previously the presence of citrullinated fibrinogen could only be detected in RA synovial fluid (41). Later research by Zhao *et al.* (2008) confirmed the presence of citrullinated fibrinogen containing immune complexes in RA plasma (131). The insolubility of fibrin may increase the likelihood of it being citrullinated in circulation. As we could not stain specifically for citrullinated residues in fibrin only, detected fluorescence could also have originated from citrullinated histone derived from neutrophil extracellular traps (135). Nevertheless, binding of ACPAs to fibrin could render it less degradable, by decreasing available binding surface to plasmin (136). There remains conjecture as to the effect of citrullination on hemostatic outcome. Citrullination of proteins results in structural unfolding (42) and loss of function (137), which increases its antigenic shelf-life. It has been demonstrated that citrullinated fibrinogen is resistant to thrombin digestion, as preferential epitopes for PADs overlap with thrombin binding sites (45, 46, 138). Despite this, fibrinogenesis in RA is by no means impaired, as evidenced by this study and others. It is plausible that high levels of fibrinogen (10) and thrombin activity (17, 139) in RA has much stronger influence on the fate of fibrinogen than PAD enzymes. There is also evidence that upstream coagulation factors and fibrinolytic components are susceptible to citrullination (140, 141). It is therefore difficult to predict a hemostatic endpoint based on overall citrullination and the effect of citrullination on thrombosis cannot be postulated on singular reactions. The implications that citrullination could have on fibrin, being the end-product of coagulation and a major determinant of thrombotic risk, remains intriguing and should be further investigated.

Future investigations would be to determine the effect of citrullination on the thrombotic potential of formed thrombi. Our research has previously implicated the role of amyloid resembling structures in fibrin clots that confer a prothrombotic phenotype. There is evidence to suggest that the processes of citrullination and amyloidogenesis may overlap. Citrullination has been shown to affect the aggregation and oligomerization of β-amyloid proteins (142) and that citrullination of human myelin oligodendrocyte glycoprotein (MOG) can lead to amyloid-like behavior shift that has pathogenic implications for multiple sclerosis (143).

This study did present some limitations and challenges. The relatively small sample population complicates the determination of correlative associations between inflammatory and hemostatic indices. The determination of true overall fibrin diameter was also not possible with current techniques – statistical analysis revealed a measurement accuracy of 82%. The detection of citrullination in fibrin networks using fluorescent techniques were unspecific to fibrin(ogen) in this study. However, the current study included this analysis only as a preliminary probe into determining if citrullination of clots was a discerning factor between RA and non-RA individuals.

## Concluding remarks

Inflammatory and thrombotic processes are highly pertinent to the development of joint disease and cardiovascular complications. There is a need to study changes that occur within synovial environments in unison with simultaneously occurring changes within circulatory tracts. Further investigation into overlapping processes that are crucially involved in the concurrent development of both RA and CVD could reveal improved global disease markers and novel targets for therapeutic intervention. The formation and structure of fibrin clots in RA shows an atypical pattern compared to conventional observations of hypercoagulation under inflammatory conditions. We propose determining if citrullination causes a structural and functional shift in the nature of fibrin to represent an amyloid-like state. This protein modification could potentially contribute to the formation of aberrant fibrin clots in RA patients that confer a higher degree of thrombotic risk.

## DECLARATIONS

### Consent for publication

All authors approved submission of the paper.

## Raw data sharing

All raw data is available at: https://1drv.ms/u/s!AgoCOmY3bkKHiqIaVlNkRzHB8cUo6Q?e=FusgPo

## FIGURE AND LEGENDS

**Figure 8:** Overview of the overlapping processes of inflammation and coagulation in both synovial and vascular compartments. **1.** The chronic and systemic nature of the inflammatory response in RA characterizes the disease as an independent risk factor for CVD. **2.** The movement of leukocytes, inflammatory cytokines, procoagulant factors and immune complexes are aided by vascular endothelial dysfunction and neovascularization of hyperproliferative joint tissues. **3.** The role of fibrin(ogen) is integral to the formation of hyperplastic and destructive synovial tissue (pannus) and vascular thrombosis, while being a prominent self-protein target of aberrant citrullination and autoimmunogenicity in RA.

**Figure 9:** Fibrin fiber diameter measurement scheme. SEM micrographs of a prepared PPP clot from a representative healthy control (top) and RA (bottom).

**Figure 10:** Box-and-whisker plots for statistically significant TEG^®^ and V-plex parameters. Boxes represent the median and IQR. Whiskers indicate upper (75^th^ percentile + 1.5*IQR) and lower (25^th^ percentile −1.5*IQR) extremes. Outlier values are indicated by black dots

**Figure 11:** Lattice of cross-plots and correlation values across statistically significant TEG^®^ and V-Plex measurements. The sample values are indicated by crosses (RA: red, Control: black). Strong intra-analysis correlation exists between CRP with SAA and sICAM-1, between sICAM-1 and VCAM-1, and between all TEG parameters (R, K, Angle, TMRTG). CRP and SAA show moderate correlation with inflammatory and endothelial markers.

**Figure 12:** Scanning electron micrographs of the fibrin network ultrastructure. Representative micrographs of the fibrin network in healthy controls (A) and RA patients (B-D). The altered clot ultrastructure in RA, consisting of less permeable networks of thicker fibrin fibers, represents a prothrombotic phenotype.

**Figure 13:** Confocal microscopy of PPP clots stained with a Citrulline monoclonal antibody. Confocal micrographs of representative control and RA samples. Each row represents identical areas captured of fibrin clot preparations using transmitted light to illustrate fibrin network topography (left-hand column) and a green channel (middle column) to identify fluorescent particles corresponding to citrulline residues. Fluorescent images are also provided in binary (right-hand column) to better illustrate differences seen between RA and control samples with respect to citrulline-coupled fluorescence. Particle analysis confirmed the observable presence of enhanced fluorescent signal in RA samples (n=10) versus healthy controls (n=10).

**Figure 14:** Box-and-whisker plots for micrographs analysis. Outlier values are indicated by black dots. In comparison to healthy individuals, RA subjects exhibited significantly increased fibrin fiber diameters (median: 214nm vs 120nm, OR=22.7) and detected fluorescent areas (median: 3279 vs 1095, OR=2.23) of fibrin citrullination.

**Table 5:** Demographic and clinical characteristics of study participants.

**Table 6:** Vascular injury panel (V-Plex) analysis.

**Table 7:** Thromboelastography^®^ analysis.

**Table 8:**Microscopy Image Analysis.

